# Enhancing the outcome of crystallographic screening for efficient drug discovery by choosing the right crystal form

**DOI:** 10.1101/2025.08.20.671191

**Authors:** Tatjana Barthel, Jan Wollenhaupt, Laila S. Benz, Patrick Reinke, Linlin Zhang, Melanie Oelker, Frank Lennartz, Helena Tabermann, Uwe Mueller, Alke Meents, Rolf Hilgenfeld, Manfred S. Weiss

**Author notes:** Shared first authors (Tatjana Barthel, Jan Wollenhaupt).

## Abstract

In more and more drug discovery projects, crystallographic fragment screening (CFS) is employed as an early screening method. Here, we demonstrate that choosing the right crystal form has a profound influence on the hit rates and hence success and speed of downstream lead generation. Two CFS campaigns with the same fragment library and an almost identical experimental setup were carried out against the two crystal forms of the SARS-CoV-2 main protease.While both crystal forms exhibit similar diffraction properties, the observed hit rates in the two campaigns were vastly different. For the monoclinic crystals a hit rate of 3% was determined, while a hit rate of 16% was observed for the orthorhombic crystals. These findings align with the more open molecular packing in the orthorhombic crystals where the solvent channels leading to the active sites are about twice larger than in the monoclinic crystal form. Our results highlight the critical importance of the crystal system in a crystallographic fragment-screening campaign and identify this parameter as one of the most important ones to be optimized during preparation of a campaign.

## Introduction

Within the past five years, the SARS-CoV-2 pandemic has caused over 778 million infections and over 7 million deaths in the last five years (https://covid19.who.int/, 20.08.2025). While the rapid development of several mRNA vaccines has helped to reduce the number of severe disease progressions, antiviral compounds still play a key role in helping those who are already infected, in particular with more and more immune escape virus variants on the horizon (Lauring *et al*., 2022; Ao *et al*., 2023). One of the few drugs being administered against SARS-CoV-2 is Paxlovid, which contains the active site inhibitor of the main protease of SARS-CoV-2 (MPro) nirmatrelvir (Owen *et al*., 2021; Tuttle *et al*., 2025). MPro is responsible for virus maturation by processing the viral polypeptide chain at eleven sites, including its own N-and C-terminus. Since MPro is essential for the viral life cycle, inhibiting the protease stops the virus from replicating. MPro is a particularly promising target because there are no human orthologues of the protease, reducing the risk of off-target effects (Zumla *et al*., 2016). Although MPro has a relatively low mutation rate, increased selection pressure from nirmatrelvir and natural mutations have already led to the emergence of nirmatrelvir-resistant MPro variants, as shown in recent reports (Tamura *et al*., 2024; Yamamoto *et al*., 2024; Zuckerman *et al*., 2024). This highlights the urgent need for new potent anti-SARS-CoV-2 drugs, as well as preparedness strategies for future pandemics involving related coronaviruses.

Especially in cases like the SARS-CoV-2 pandemic, but also in general, efficient drug discovery is essential. In the last decades fragment screening was established as a beneficial method to lower costs and increase chemical diversity (Li, 2020; Xu & Kang, 2025; Erlanson et al., 2016). Crystallographic fragment screening (CFS) has been shown to be an effective method Crystallographic fragment screening (CFS) has been shown to be an effective method to find initial starting points for drug development and allows the rapid design of follow-up compounds by structure-based design (Schiebel *et al*., 2016; Bentley *et al*., 2020; Metz *et al*., 2021). CFS was also applied intensely in the last five years for the search of potent SARS-CoV-2 antivirals, the major example being the COVID Moonshot consortium (COVID Moonshot Consortium *et al*., 2021). Within the consortium, over 150 active participants from different technological backgrounds combined their expertise and were able to generate huge amounts of open-source data towards an anti-SARS-CoV-2 drug. Over 18,000 compounds were designed, over 840 crystallographic data sets of bound compounds were collected and over 2,400 novel compounds were synthesized. This effort has resulted in a candidate for clinical trials (Boby and Fearon *et al*., 2023), providing a great example of an open and collaborative scientific effort and highlighting the benefits of CFS.

For a successful screening campaign, the crystal packing of the protein must ensure the accessibility of the sites of interest via the solvent channels. Large solvent channels and accessible binding sites are highly preferred, if not indispensable for success. In order to obtain an optimal crystal system, it may be advantageous to test different crystal forms of the target protein (Bauman *et al*., 2013).

For MPro, which is the target of this study, two crystal forms have been reported, a monoclinic and an orthorhombic crystal form (Zhang *et al*., 2020; Noske *et al*., 2021). Most CFS campaigns against MPro have been carried out with crystals in the monoclinic space group (Douangamath *et al*., 2020; Günther *et al*., 2021), although it has been discussed in the literature that the more open orthorhombic space group may have advantages for crystallographic and virtual screening (Costanzi *et al*., 2021; Noske *et al*., 2021; Huang *et al*., 2024). Several fragment libraries have been applied to MPro, with hit rates typically not exceeding 5% (Douangamath *et al*., 2020). This is relatively low compared to the capabilities of CFS yielding 20-30% hit rates (Wollenhaupt *et al*., 2020) or even over 40% for certain targets (Füsser *et al*., 2023). A CFS campaign against the orthorhombic crystal form had been performed beforehand though did not yield the expected increase in hit rate (Noske *et al*., 2021). As all screenings were done with various parameters differing in the experimental set up, it is unclear how much the crystal packing played a role.

Here, we present the first direct comparison of two crystallographic fragment screening campaigns using the F2X-Entry Screen (Wollenhaupt *et al*., 2020) against the two reported MPro crystal forms. In line with previous results, we achieved a relatively low hit rate of 3% for the monoclinic system but could obtain a significantly higher success rate of 16% for the orthorhombic system. This observation is vital for future fragment-based drug design projects to reach high-quality starting points more efficiently.

## Materials & Methods

### Protein Expression and Purification

The MPro gene was cloned into a pGEX-6-1 vector with a C-terminal His_6_-tag. (Zhang *et al*., 2020). The plasmid (provided by Lin Zhang from the Hilgenfeld group) was transformed into *E. coli* strain BL21-Star (DE3). An overnight culture was inoculated from transformed clones and transferred the next day into auto-induction medium (Studier, 2005). The cells were grown at 37°C until OD=0.6 and then the temperature was lowered to 18°C for overnight expression. Cells were harvested the next day through centrifugation at 10,000g for 10 min. Cell pellets were flash-frozen in liquid nitrogen until purification. For purification, the cells were thawed and resuspended in 20 mM Tris pH 7.8, 150 mM NaCl, 5 mM Imidazole, 0.05% (v/v) Tween. The cells were lysed via sonication and centrifuged at 40.000g for 45 min at 4°C. The supernatant was loaded onto a Ni-NTA affinity chromatography column, and the protein was eluted with a stepwise imidazole gradient up to 500 mM imidazole. The protein was dialyzed overnight with addition of 1:10 w/w PreScission into 20 mM Tris pH 7.8, 150 mM NaCl, 1 mM DTT. The next day the cleaved protein was subjected to a Ni-NTA affinity chromatography column to remove the cleaved His_6_-tag. The final purification step was size exclusion chromatography using a Superdex 75 column equilibrated in 20 mM Tris pH 7.8, 150 mM NaCl, 1 mM TCEP, 1 mM EDTA. The fractions with the purified protein were pooled, concentrated to 5 mg/ml and flash-frozen in liquid nitrogen before storage at −80°C.

### Crystallization

MPro was crystallized in the orthorhombic space group in 23.5% (w/v) PEG 1.500, 0.2 M MIB pH 7.7, 5% (v/v) DMSO, 1 mM TCEP and 0.025 mM EDTA pH 7.0 using the NT8 pipetting robot (Formulatrix) and MRC 3-lens 96-well low-profile plates. The final drop consisted of 200 nl protein, 100 nl reservoir and 50 nl 1:50 seed stock dilution and was equilibrated at 20°C against 40 µl reservoir. Initial seeds for the orthorhombic space group were kindly provided by Deniz Eris (Macromolecular Crystallography group, Paul Scherrer Institute, Villigen, Switzerland). Based on the crystals grown from the initial seeds, new seeds were prepared in the following way. Crystals from one drop were crushed and transferred into 50 µl reservoir solution. The mixture was vortexed four times for 30 s with 30 s on ice in between with a Seed Bead™ (Hampton Research). The final seed stock was diluted to 1:50, aliquoted and flash-frozen in liquid nitrogen. Crystals grew within 2 days.

Protein crystals grown in the monoclinic space group were produced as described before in Günther *et al*., 2021.

### Crystallographic Fragment Screening – Soaking and Data Collection

The F2X-Entry Screen was screened against MPro in both space groups. For both campaigns the following soaking buffer was used: 23.5% (w/v) PEG 1.500, 0.2 M MIB pH 7.7, 5% (v/v) DMSO, 1 mM DTT and 0.025 mM EDTA pH 7.0. A soaking plate was prepared with 40 µl crystallization solution as reservoir and a 0.4 µl drop of the respective soaking solution onto the dried-on fragments. The crystals were transferred from the crystallization plate into the soaking drops. After the transfer the plate was sealed with crystallization foil and incubated overnight at 20°C. The next day the crystals from each soaking drop were harvested, flash-cooled in liquid nitrogen and stored until measurement. Each fragment soak was named according to the fragment’s placement on the F2X-Entry Screen plate (A01 – H12). Due to the collection of duplicates, the samples also received indicators to achieve unique names for each sample/dataset (A01a, A01b – H12a, H12b).

Data collection was performed at BL14.1 (HZB BESSY II, Berlin, Germany).(Mueller *et al*., 2025) Diffraction data from crystals belonging to the monoclinic space group was collected with 1500 images in 0.2° increments, an exposure time of 0.1 s, a 100 µm aperture, detector distance corresponding to 1.35 Å maximum resolution at the detector edge and at an X-ray energy of 13.5 keV. The data of the orthorhombic space group CFS campaign was collected with 800 images in 0.2° increments, an exposure time of 0.1 s, a 100 µm aperture, detector distance to 1.35 Å maximum resolution at the detector edge and at an energy of 13.5 keV.

### Crystallographic Fragment Screening – Data Analysis

The collected data was then subjected to FragMAXapp (Lima *et al*., 2021) and processed automatically via XDSAPP (Sparta *et al*., 2016) and afterwards automatically refined with fspipeline. (Schiebel *et al*., 2016) The PDB structure 6Y2E (Zhang *et al*., 2020) for the monoclinic space group C2 and 7BB2 (Costanzi *et al*., 2021) for the orthorhombic space group P2_1_2_1_2_1_ crystals were used as an input model for molecular replacement. PanDDA (Pearce *et al*., 2017) was performed for both campaigns and fragments were fitted into the event maps. In case of the orthorhombic space group CFS campaign clustering was performed via cluster4x (Ginn, 2020). By clustering the data with cluster4x three separate clusters were identified. To run PanDDA for each cluster, but at the same time consider all available datasets, PanDDA was run in a specialized way. For each cluster a PanDDA run was performed, where only the datasets associated with one cluster were used for the ground state calculation. Each fragment-bound structure of both space groups was subjected to manual building in pandda.inspect using the PanDDA event maps as the main readout. Afterwards the data was exported via pandda.export and the ensembles were split into their bound and ground state via the giant.split_conformations script. The bound state models were built with the help of the PanDDa event and Z map using Coot (Emsley *et al*., 2010). The ground state models for each bound fragment were built with the help of the average map calculated by PanDDA for the respective cluster run. The final models were merged again via the giant.merge_conformations script and then a pandda.quick_refine script was applied for the final refinement of the ensemble. All refined data was deposited in the PDB with the following IDs: monoclinic space group: group deposition ID G_1002326, 7HUC, 7HUD, 7HUE, orthorhombic space group: group deposition ID G_1002335, 7I13, 7I14, 7I15, 7I16, 7I17, 7I18, 7I19, 7I1A, 7I1C, 7I1D, 7I1E, 7I1F, 7I1G, 7I1H, 7I1I, 7I1J. Table1 for each dataset is given as supplementary data.

## Results

### Comparison of the two space groups for MPro crystal

For MPro two crystal systems have been reported: a monoclinic system of space group C2 and an orthorhombic system of space group P2_1_2_1_2_1_. The respective crystal metrics as well as the unit cell content are given in Table 1. A comparison of the molecular packing in both crystal forms reveals the vastly differing accessibility of the MPro active site (**Figure 1**). In the monoclinic space group, the active site is partially obstructed, leaving little room for compounds to bind and side chains to move. In the orthorhombic space group, the active site is much more accessible, allowing compounds to diffuse into the active site more easily and side chains to accommodate compounds binding.

**Figure 1:**
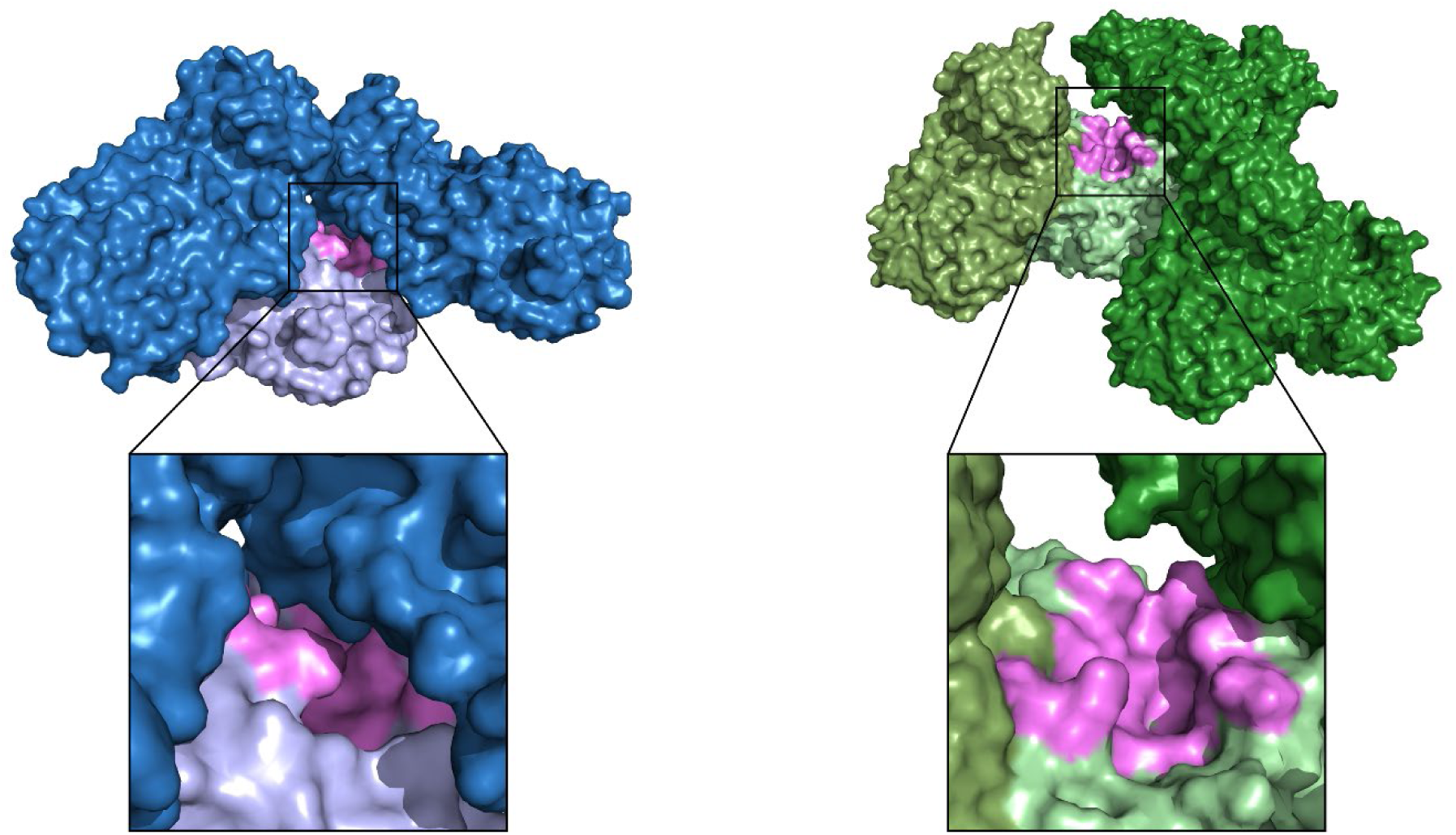
Comparison of the crystal packing and active site accessibility in the two crystal forms of MPro. The protein structures for the monoclinic space group C2 are shown in blue colors. The protein structures for the orthorhombic space group P212121 are shown in green colors. The monomer structure of MPro is presented always in a light color, while the crystal mates are shown in darker colors. In case of the orthorhombic space group, the dimer is present in the asymmetric unit, therefore a third color tone was chosen to indicate the second monomer. The active site is colored in both cases pink. A zoom in into the active site is given, highlighting the accessibility of the active site in both crystal forms. The same color scheme (blue for C2 structures and green for P212121 structures) is used throughout the manuscript.

**Table 1:** Comparison of the two MPro space groups. The values have been taken from the datasets of 6Y2E (monoclinic, Zhang *et al*., 2020) and 7BB2 (orthorhombic, (Costanzi *et al*., 2021)). The solvent content and V_M_ (Matthews Coefficient) value were calculated by Phenix Xtriage. (Liebschner *et al*., 2019).

|  | Monoclinic form | Orthorhombic form |
| --- | --- | --- |
| **Space group** | C2 | P2<sub>1</sub>2<sub>1</sub>2<sub>1</sub> |
| **Cell parameters (a, b, c [Å],**<br>**α, β, γ [°])** | 113.57, 53.75, 44.66,<br>90, 101.64, 90 | 67.84, 99.55, 103.68<br>90, 90, 90 |
| **No. of molecules per asymmetric unit** | 1 | 2 |
| **Solvent content [%]** | 38 | 52 |
| **V<sub>M</sub> (Matthews Coefficient [Å<sup>3</sup>/Da])** | 2.0 | 2.5 |

This comparison raises the question of which of the two crystal systems is more suitable for a CFS campaign. Assessing this suitability involves the rather time-consuming characterization of the diameter of each of the available solvent channels in the crystal. Also, the accessibility of the active site needs to be checked, which requires a closer look at dynamic loops and residues, nearby symmetry mates and other molecules binding to the active site. Using the recently published tool LifeSoaks (Pletzer-Zelgert *et al*., 2023), the size and shape of crystal solvent channels as well as their constriction sites can be analyzed easily automatically. Since LifeSoaks was not yet available at the time of the here described CFS campaigns, a LifeSoaks analysis was carried out in retrospect. LifeSoaks was applied to one ligand-bound dataset from each of the two CFS campaigns. As shown in **Table 2**, the orthorhombic P2_1_2_1_2_1_ crystal form has an overall bottleneck radius more than twice as large than the monoclinic C2 crystal form, and the inner and outer active site radius is 1.7 and 3.3 times larger, respectively. These differences are likely the cause of the observed higher success rate using the orthorhombic crystals. Furthermore, the difference in active site openness could potentially facilitate subsequent fragment growing or merging.

**Table 2.** LifeSoaks values for both crystal systems, calculated based on a representative fragment-bound structure identified during this study. LifeSoaks was used with the default values for the minimal channel radius of 1.7 Å and with the unified atom radius of 1.7 Å for each atom in the model. For both analysis the bound ligand was used to generate a binding pocket and then used as a reference ligand.

|  | **C2 ligand B08** | **P2<sub>1</sub>2<sub>1</sub>2<sub>1</sub> ligand G09** |
| --- | --- | --- |
| **Overall bottleneck radius [Å]** | 3.6 | 7.6 |
| **V<sub>M</sub>(Matthews Coefficient) [Å<sup>3</sup>/Da]** | 1.9 | 2.6 |
| **Inner active site radius [Å]** | 2.3 | 3.9 |
| **Outer active site radius [Å]** | 2.3 | 7.6 |

Many of the initial crystallization conditions reported for the P2_1_2_1_2_1_ crystal form in the PDB could not be reproduced in our lab or yielded again crystals in the C2 crystal form (data not shown). It turned out to be necessary to add P2_1_2_1_2_1_ seeds (kindly provided by Paul Scherrer Institute colleagues) to the initial crystallization to enable the growth of the orthorhombic crystals. Both crystal systems were then optimized for the CFS soaking process. The F2X-Entry Screen fragment library (Wollenhaupt *et al*., 2020) was used for screening. **Figure 2** shows an overview of the main quality indicators for the campaigns. Table 1 for each dataset with a fragment bound is given in the supplementary data as **Table S1**.

**Figure 2:**
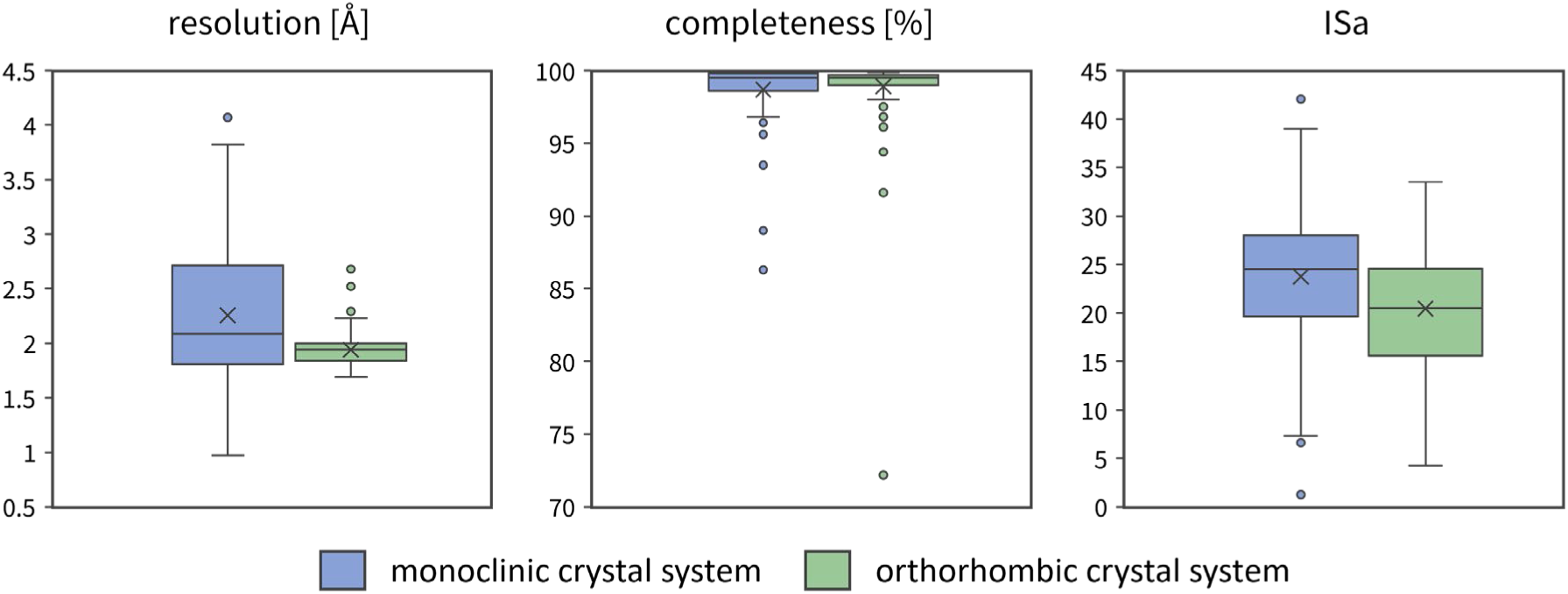
Data quality indicators for both crystal forms (monoclinic in blue and orthorhombic in green) shown as boxplots. Each boxplot shows the 25th and 75th percentile as a box, with the median as a line in the middle. Whiskers show the range of data within 1.5-fold interquartile range. Data outside this range are considered outliers and depicted as dots. The boxplots were made using Microsoft Office Excel (Version 2507).

The C2 campaign exhibits slightly lower median resolution (2.19 Å) and a larger range (1.33–3.87 Å) compared to the P2_1_2_1_2_1_ campaign (median 1.94 Å, range 1.69–2.68 Å). The completeness is on average similar in both campaigns with median completeness over 98% in both campaigns. The ISa values of the datasets from the C2 campaign were slightly higher (median 24) than for the P2_1_2_1_2_1_ campaign (median 20). Overall, both campaigns show a similar data quality based on the metrics presented.

### Fragment hits identified from monoclinic C2 screening

For the C2 campaign, three fragment hits and four binding events were identified. This translates to a hit rate of 3%. Fragments B08 and D08 bound to the active site, while fragment B07 yielded two binding events: one at the active site and one at a remote binding site. **Figure 3** shows an overview of the binders.

**Figure 3:**
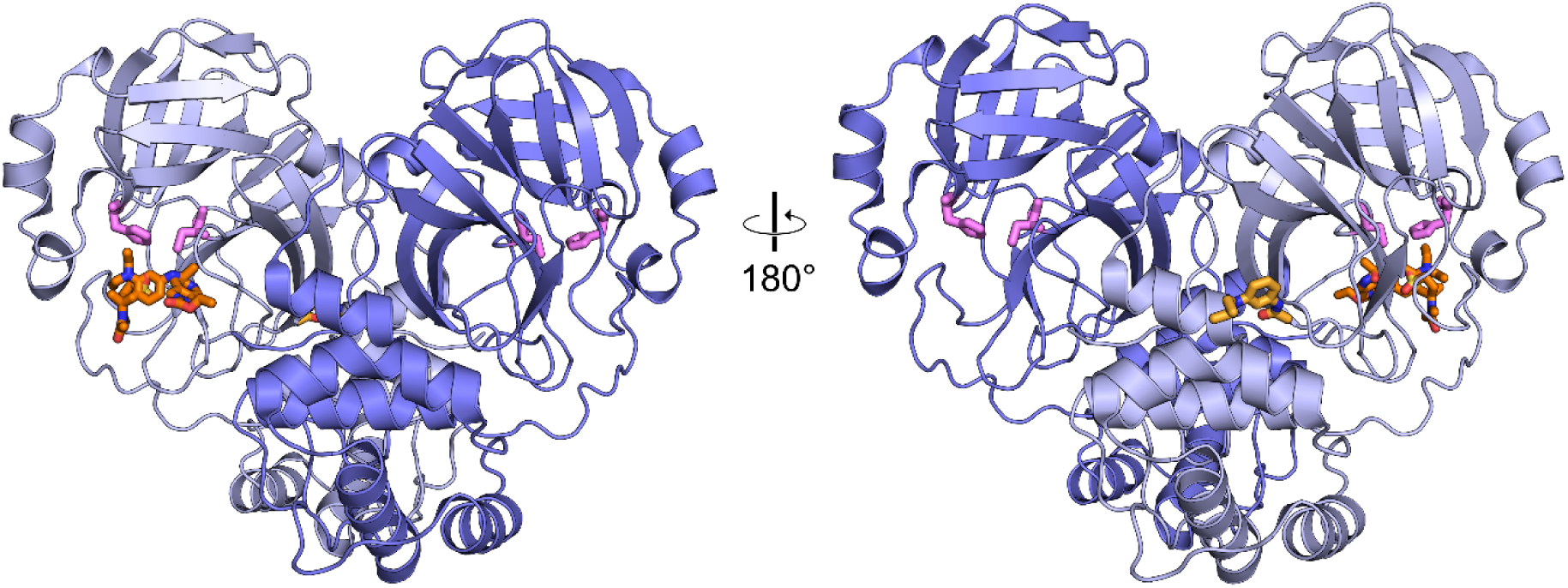
Overview of hits identified in the CFS campaign of the monoclinic C2 crystal form. The monomer of the asymmetric unit is shown in dark blue and a symmetry related protein molecule in light blue, to illustrate the biological dimeric form of MPro. The bound fragment hits are presented as orange sticks. The two catalytic side chains His41 and Cys145 are depicted as spheres colored pink. The fragments bound are only shown for the monomer of the asymmetric unit, The front and back view are depicted.

The active site binders overlap marginally and bind in different specificity pockets of MPro (**Figure 4A**). Compound B08 binds to S1, compound D08 binds to S2 and S4 and compound B07 bridges from S1 to S2 also covering S3. The overlapping structural features of the active site binders do not show similar interactions in the specificity pockets (**Figures 4B,C,E-H**). The remote binding site of B07 is close to a crystal contact (**Figure 4D**).

**Figure 4:**
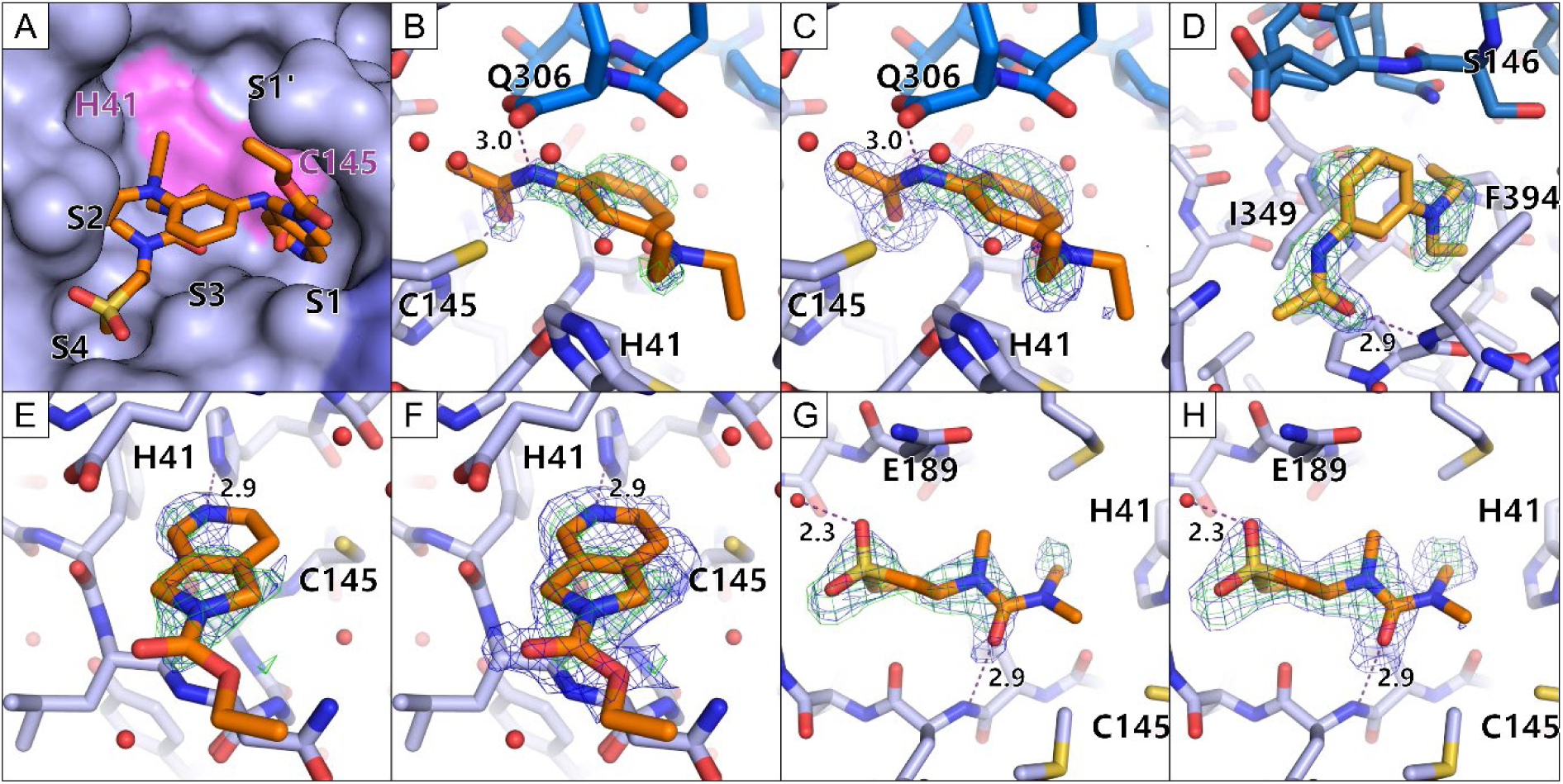
Enlarged views of the bound fragment hits in the monoclinic crystal form. The fragments are colored orange, the active site binders in a darker orange and the remote binders in a light orange. Water molecules are depicted as red spheres. Hydrogen bonds are depicted as purple dashed lines with their respective length written beside the line. The electron density is presented for the fragments (PanDDA event map is shown in blue (σ=2 for B, D, E and G; σ=1 for C, F and H) and the PanDDA Z map is shown in green (σ=3). A) A zoom in view of the active site with indicated specificity pocket positions. The protein is shown in a surface view while the fragments are shown as sticks. The active site His41 and Cys145 are shown in pink. B and C) The fragment B07 is shown with different sigma levels for the PanDDA event map for clear interpretation. The C-terminus of the second monomer, making up the dimer is shown in darker blue. D) The second binding site of fragment B07 is shown at a remote binding site close to a crystal contact. E and F) The fragment B08 is shown with different sigma levels for the PanDDA event map for clear interpretation. G and H) The fragment D08 is shown with different sigma levels for the PanDDA event map for clear interpretation.

### Fragment hits identified from orthorhombic P2_1_2_1_2_1_ screening

For the P2_1_2_1_2_1_ campaign 11 fragment hits could be identified binding to the protein. Out of these fragment hits two bound at remote binding sites (G03, H03) and 7 fragments bound at the active site (B05, B08, C10, D04, D08, F04, G09). One fragment hit bound twice to the protein, once in the active site and once at a remote binding site (C02). One fragment hit bound three times to the protein, twice in the active site and once at a remote site (D11). This amounts to 14 binding events overall and a hit rate of 10%. The analysis of this dataset was further improved using cluster4x for picking more homogenous datasets for the analysis. This step had been shown before to increase homogeneity of the dataset which is crucial for a PanDDA-based hit identification (Barthel *et al*., 2022; Ginn, 2020). Clustering of the dataset resulted in three distinct clusters. For each of the clusters, a separate PanDDA run was performed. The application of this step resulted in five additional fragment hits. Taken together, this campaign resulted in 16 fragments overall, 19 binding events and a hit rate of 16%. Out of the 16 fragment hits, three fragments bound at remote sites (G03, H03, plus H11). 11 fragments bound at the active site (B05, B08, C10, D04, D08, F04, G09, plus C07, E11, G04, G10). One fragment bound twice, once at the active site and once at a remote binding site (C02). One fragment hit bound three times to the protein, twice in the active site, but with different binding poses in the two active sites present in the asymmetric unit and once at a remote site (D11). An overview of the binders is depicted in **Figure 5**.

**Figure 5:**
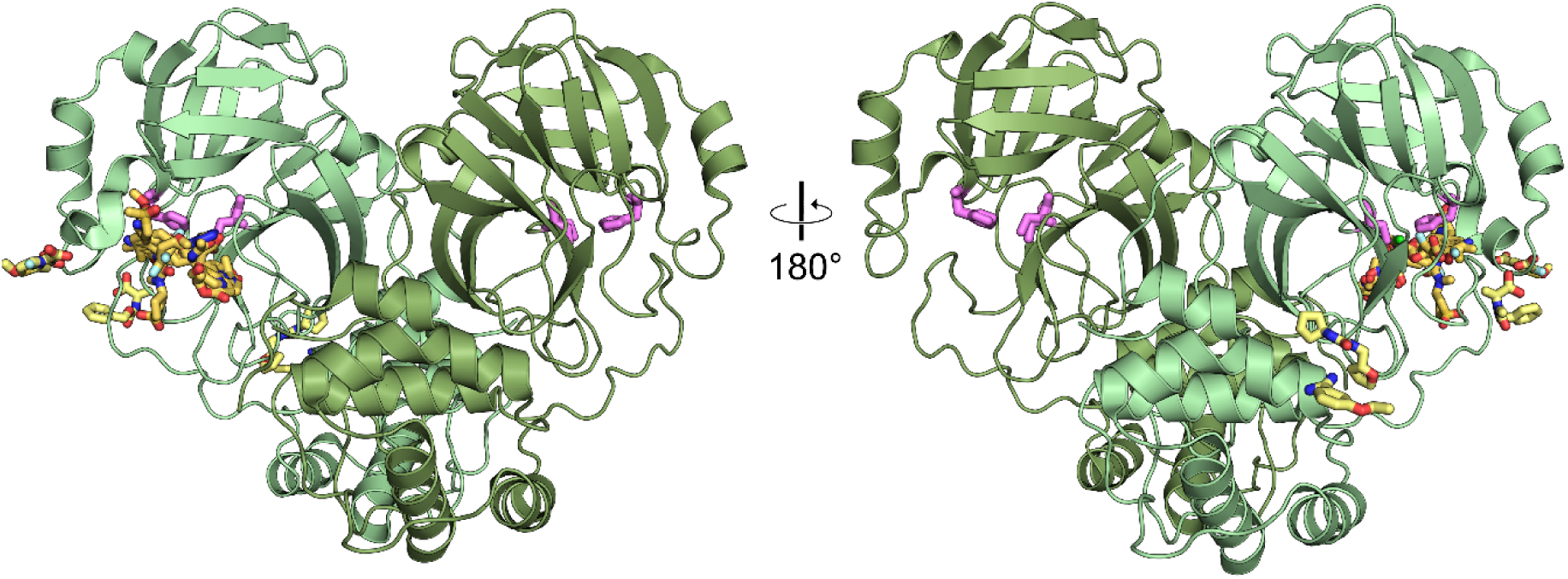
Overview of hits identified in the CFS campaign of the P2_1_2_1_2_1_ crystal form. The monomers are shown in green colors as cartoon and the bound fragment hits are presented as yellow sticks. The two catalytic side chains His41 and Cys145 are depicted as sticks colored pink. The fragments bound are shown in one monomer’s active site as the binding mode was similar in both active sites. The front and back view are depicted.

The active site binders cover the specificity pockets S1, S2, S3 and S4 (**Figures 6A, B**). The fragments B05, B08, C07, D11 and E11 bind in the S1 pocket (**Figures 6C, D, G, K, L, N**). G09 reaches over several pockets from between S1’ and S1 to S3 and S2 (**Figure 6R**). G10 covers a similar area to G09 but is much shorter and does not reach deep into the pockets. G10 differs from all other fragments as it bound covalently to the active site Cys145 based on the electron density evidence (**Figure 6S**). D08 bound in S4 reaching into S2 (**Figure 6J**). C02, C10 bind mainly in S2 (**Figures 6E, H**), while D04, F04, G04 bind in S2 and reach along the active site, though not into one of the specificity pockets (**Figures 6I, O, Q**).

**Figure 6:**
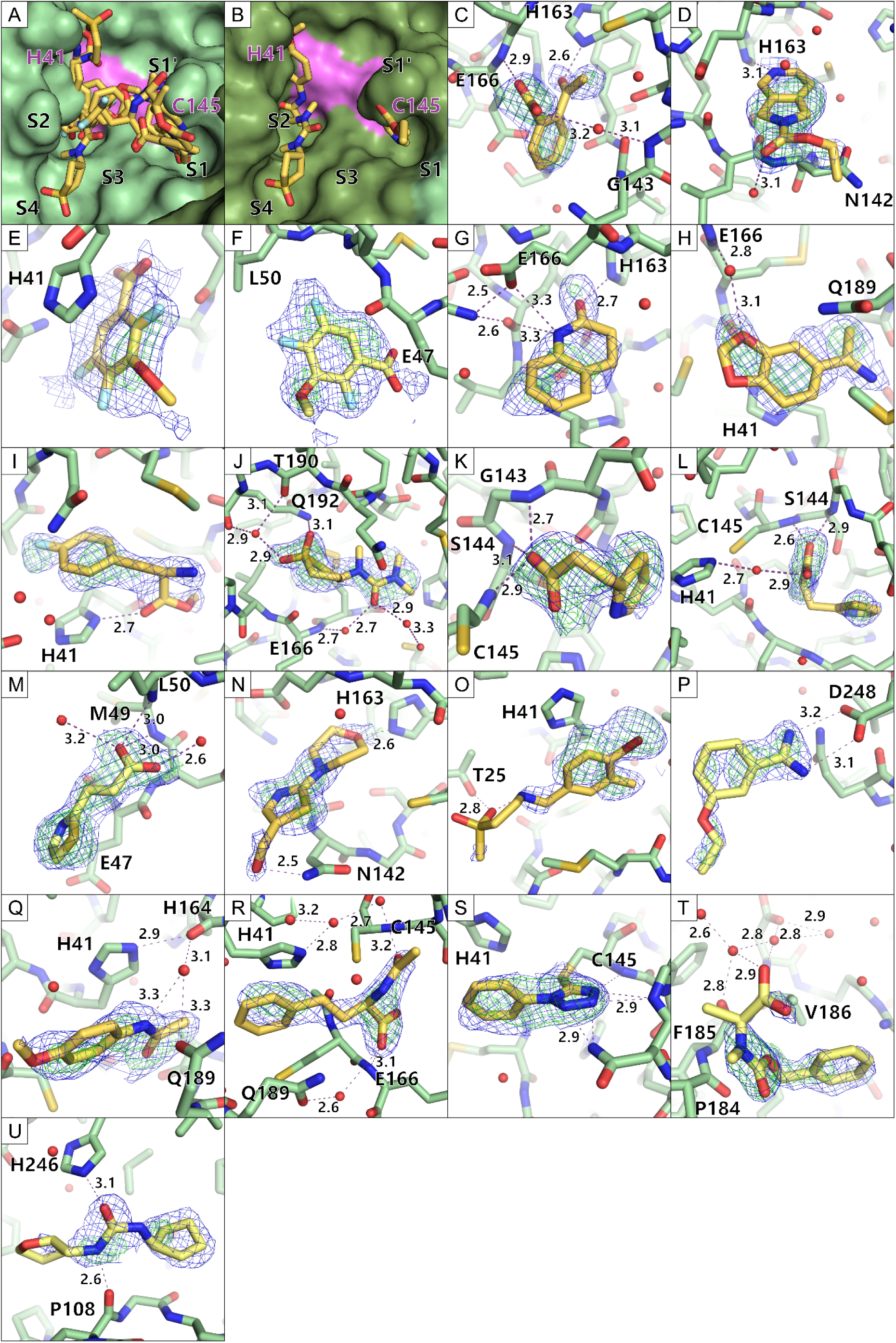
Enlarged view of the bound fragment hits in the orthorhombic crystal form. The fragments are colored in yellow, the active site binders in a darker yellow and the remote binders in a lighter yellow. Water molecules are depicted as red spheres. Hydrogen bonds are depicted as purple dashed lines with their respective length written beside the line. The electron density is presented for the fragments (PanDDA event map is shown in blue (σ=2) and the PanDDA Z map is shown in green (σ=3)). A-B) A zoom in view of the active site with indicated specificity pocket positions. The protein is shown in a surface view while the fragments are shown as sticks. The active site His41 and Cys145 are shown in pink. C-T) Each fragment bound to the protein is shown as sticks. The protein is shown as sticks too. (C=B05, D=B08, E=C02 in active site, F=C02 at remote site, G=C07, H=C10, I=D04, J=D08, K=D11 in monomer 1, L=D11 in monomer 2, M= D11 at remote site, N=E11, O=F04, P=G03, Q=G04, R=G09, S=G10, T=H03, U=H11).

The remote binders G03 and H11 bind the protein at solvent exposed surfaces (**Figures 6P, U**). These binding sites have not been seen before and merit further investigation. The second binding events of C02 and D11 remote from the active site were close to crystal contacts (**Figures 6F, M**). Additionally, H03 also bound close to a crystal contact (**Figure 6T**). These three events must be studied critically to determine if they are crystal artifacts or interesting binding sites.

### Comparison of the two campaigns

Combining both campaigns, 17 unique fragments of the F2X-Entry Screen could be identified. Fragments B08 and D08 were the only two overlapping hits. Both fragments bind in both campaigns in a similar binding pose (**Figure 7**) providing mutual pose validation. B08 bound in almost the exact binding pose (**Figures 7A, C, E**). The position of D08 differed slightly, probably due to the more constricted packing of the protein in the monoclinic crystal form. The loop containing T190 and A191 is able to move upwards as indicated by a magenta arrow (**Figures 7B, D, F**). The higher flexibility in the orthorhombic crystal form allows for the fragment to position itself in a way that a more elaborate hydrogen bond network can be established facilitated by water molecules.

**Figure 7:**
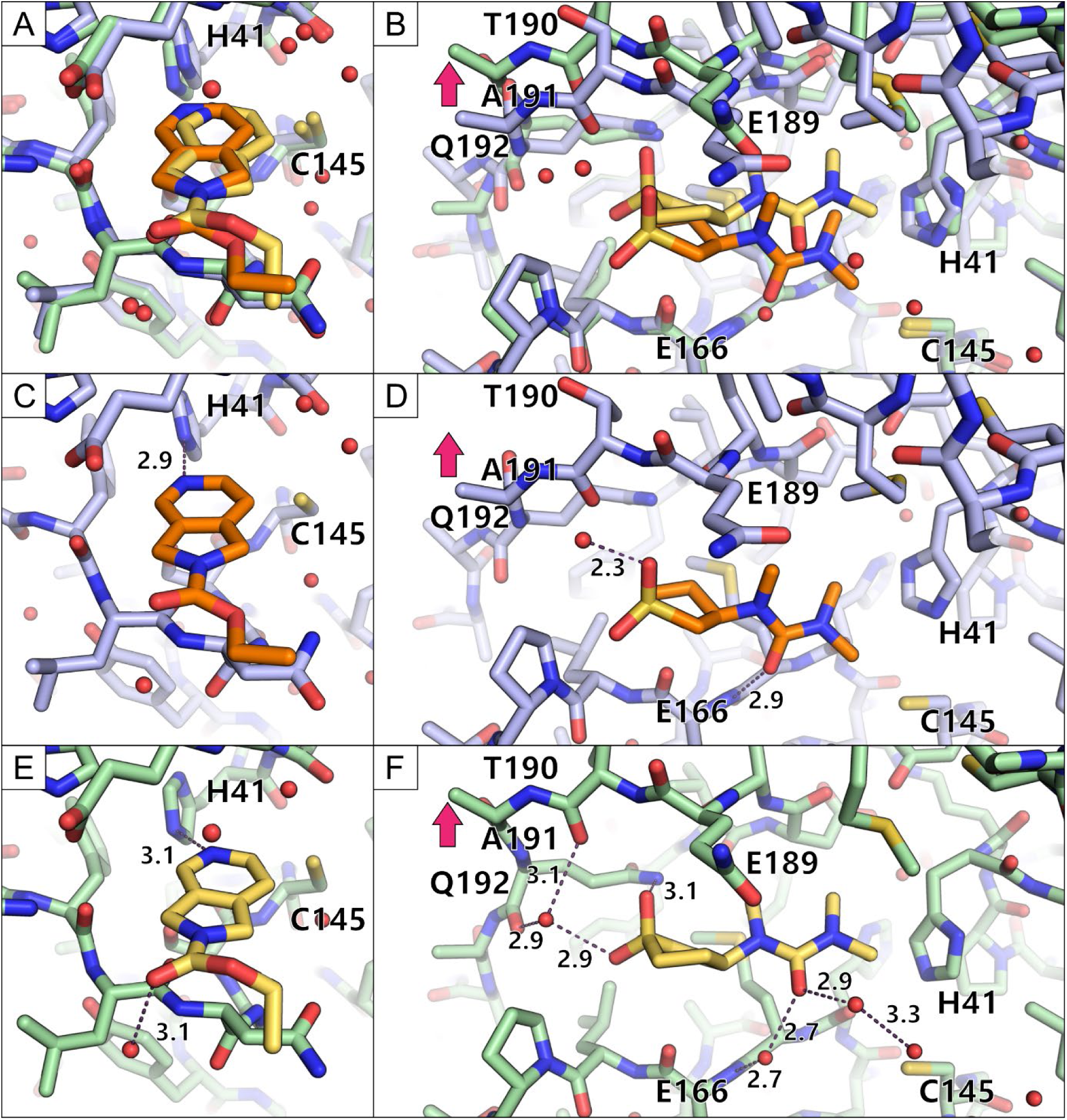
Comparison of the binding events of B08 and D08 from both campaigns. The binding pose from the monoclinic campaign is colored orange while the binding pose from the orthorhombic campaign is shown in yellow. The protein is shown in blue for the monoclinic campaign and in green for the orthorhombic campaign. Water molecules are depicted as red spheres and hydrogen bonds are shown as purple dashed lines with their distance written next to them. A-B) An overlay is shown for B08 (A) and D08 (B) of both campaigns. C-D) The binding pose of both fragments in the monoclinic space group is highlighted. E-F) The binding pose of both fragments in the orthorhombic space group is depicted.

Fragment B07 had been found to bind in the monoclinic C2 campaign but not in the orthorhombic P2_1_2_1_2_1_ campaign. Upon closer investigation of the B07 binding to the protein, additional density of the C-terminus of a symmetry related molecule became visible (**Figure 8**). Starting from the residue Ser301 the C-terminus adopts a different conformation in the bound state of B07. The C-terminus interacts with the fragment by stacking of the aromatic benzene ring with the peptide backbone and a hydrogen bond between the amide nitrogen of the fragment with the terminal carboxyl-group of Gln306. The C-terminus also engages in hydrogen bonds with N142.

**Figure 8:**
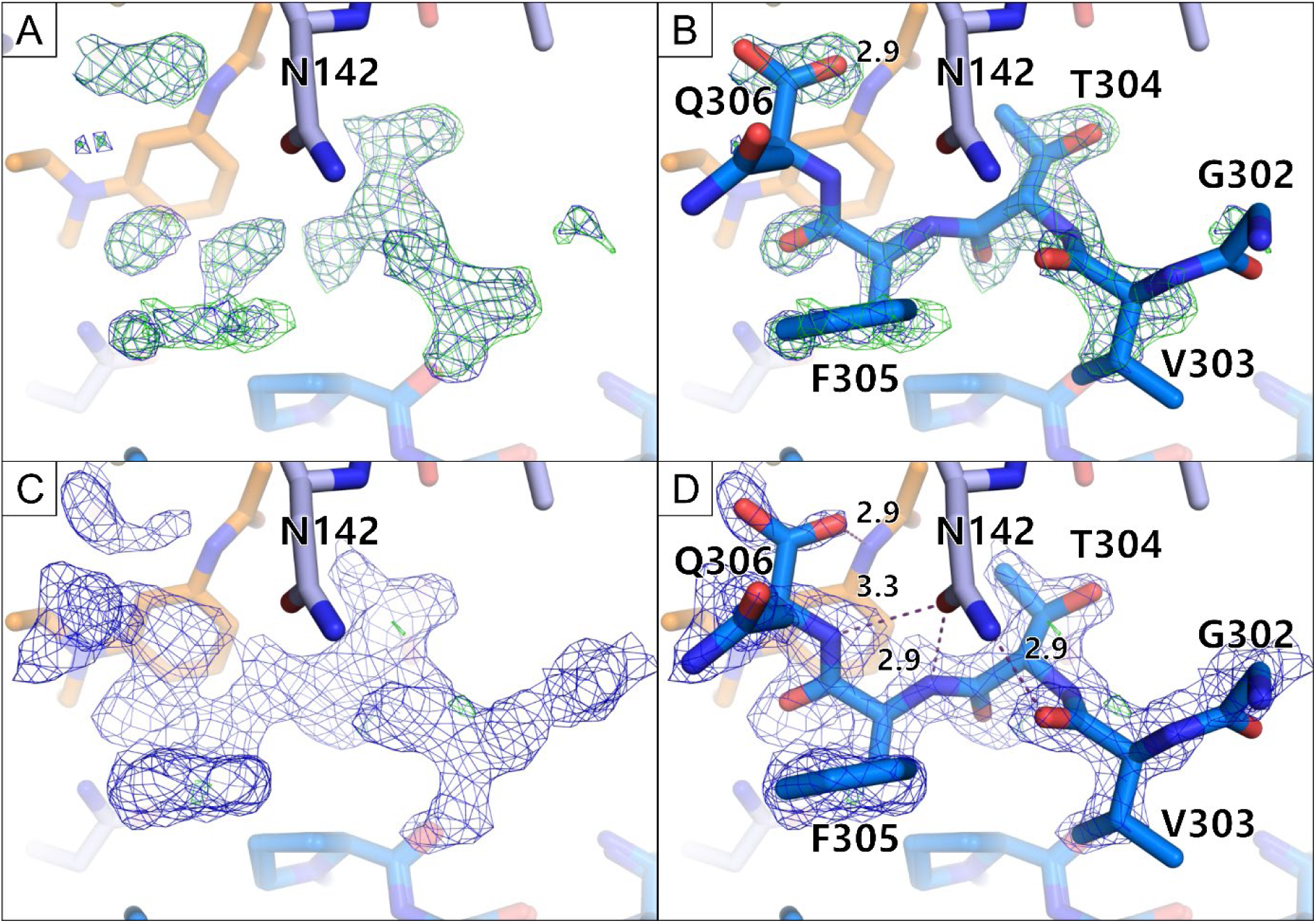
The C-terminus of a symmetry-related molecule binding to the B07 binding site. The MPro monomer of the asymmetric unit is shown in light blue sticks. A symmetry related monomer is shown in darker blue sticks. Fragment B07 is shown in orange sticks. A) The PanDDA event map is depicted in blue (σ=2) and the Z map in green (σ=3). B) The same maps are shown as in A, but with the placed C-terminus. C) The electron density after one round of refinement is shown; (2mF_o_−DF_c_, α_c_) maps contoured at σ = 1 (blue) and the (mF_o_−DF_c_, α_c_) maps contoured at σ = 3 (green). D) The same electron density is shown as in C, but with the placed C-terminus.

### Comparison to a previously performed orthorhombic CFS campaign

The F2X-Entry Screen had been previously used to perform a CFS campaign against MPro in the orthorhombic crystal system (Noske *et al*., 2021). Based on the cell metric the crystal system described by Noske et al. (2021) is the same as the one described here. However, upon a closer look, it is revealed that Noske et al. (2021) used a slightly different construct (immature MPro) as well as different crystallization conditions and slightly different soaking parameters (**Supplementary Table 2**). As a result of their campaign Noske et al. (2021) reported three fragment hits and a hit rate of only 3 %. Interestingly, the observed hits (E03, E06, G05) are different from the ones observed in either campaign here (B05, B07, B08, C02, C10, D04, D08, D11, E11, F04, G03, G04, G09, G10, H03, H11). This comparison shows that a superficial look at space group and cell metrics is not sufficient to ensure comparability. Since a CFS experiment is close to the limit of detection of binding, differences in the protein construct and experimental parameters may result in different outcomes.

## Discussion

### Comparison of the screening campaigns using the monoclinic and orthorhombic crystal form of MPro

The success of a CFS campaign depends on a well-established and optimized crystal system. This includes reproducible crystal growth, solvent tolerance of the crystals and most importantly consistent high resolution diffracting crystals. The importance of crystal packing as a success factor in CFS campaigns has been studied before. Schuller *et al*., 2021, screened another SARS-CoV-2 protein, NSP3, in two different space groups (P2 and P4_3_) and achieved a higher hit rate of 8.8% in the favored space group compared to 5.6% hit rate in the unfavored space group. Other earlier studies on the influenza A nuclease also focused on the identification of new crystal forms of the target protein because of the otherwise occluded active site (Patel *et al*., 2014; Bauman *et al*., 2013). However, this factor is still underappreciated.

Here, we present the first systematic study, keeping all parameters consistent besides the space group and crystallization condition. Both screens were conducted with the same protein construct, the same soaking conditions, using the same fragment library and a highly similar computational workflow for data analysis.

The CFS campaign against the orthorhombic crystal system resulted in an over fivefold increase in hit rate and higher coverage of the active site compared to the campaign with the monoclinic crystal form. Next to the highly consistent experimental set up, the achieved data quality of the two screens is comparable. The CFS campaign against the orthorhombic crystal form resulted in a slightly better median resolution (better by 0.25 Å), but this alone likely does not explain the significantly increased hit rate. Our experimental findings are further supported by the *in silico* analysis of the solvent channels and the active site using LifeSoaks (Pletzer-Zelgert *et al*., 2023). The analysis highlights the increased radius of the active site and the larger overall bottleneck radius for the orthorhombic crystal form.

The crystal packing not only influenced the hit rate, but also the potential relevance of the hits. For instance, the fragment B07 binding to the active site in the monoclinic MPro form is likely only a crystallization artefact, because it clearly interacts with the C-terminus of a symmetry related protein molecule. It probably did not bind in the orthorhombic campaign as the C-terminus cannot interact in the same way in the orthorhombic crystal system. The fragment would thus probably not bind to the enzyme in its native dimeric state in solution.

In conclusion, although most of the reported screening activities have been directed at the monoclinic crystal system (Consortium *et al*., 2021; Douangamath *et al*., 2020; Günther *et al*., 2021) the orthorhombic system appears to be much better suited for CFS approaches. The LifeSoaks analysis supports our experimental findings and would be a useful tool at the start of a fragment screening campaign to choose a suitable crystal form. Our findings highlight the importance of choosing the correct crystal system for crystallographic fragment-screening for maximizing the chance of success.

## Conclusion

Taken together, a systematic comparison of two CFS campaigns on two crystal forms of MPro this work underpins the importance of crystal packing for performing successful CFS campaigns. 14 fragments were found binding to the active site of MPro and can serve as starting points for new fragment-based drug design or provide novel ideas for optimized binding in the different sub pockets to improve existing compounds. In general, the presented work exemplifies the need to investigate and to optimize the crystal packing of the target protein for future crystallographic screening campaigns.

## Supporting information

Suuplemental Table 1 and 2

## Acknowledgements

We want to acknowledge the great help by Tobias Krojer (MAX IV laboratory) for the PDB submission of the fragment-bound structures. Additionally, we thank the Macromolecular Crystallography group at the Paul Scherrer Institute for sharing their MPro P2_1_2_1_2_1_ crystal seeds with us. Additionally, we also acknowledge several grants and collaborations supporting this study: BMBF Collaborative Research Project STOP-CORONA (project no. 05K20CB1), iNext Discovery, project No. 871037, funded by the Horizon 2020 program of the European Commission; the German Research Foundation (DFG) via the project NECESSITY (FE2166/1-1); the Helmholtz Gemeinschaft via the Innovation Pool projects FISCOV and FISVIR and the Joint Berlin MX Laboratory.

