## Supplementary material for "Enhancing the outcome of crystallographic screening for efficient drug discovery by choosing the right crystal form": Suuplemental Table 1 and 2

**Orthorhombic coronavirus main protease crystals provide a higher success rate in fragment screening**

** shared first authors*

^1)^ Macromolecular Crystallography, Helmholtz-Zentrum Berlin, Albert-Einstein-Str. 15, 12489 Berlin, Germany

^2)^ present address: Proteros biostructures GmbH, Bunsenstr. 7a, 82152 Planegg-Martinsried, Germany

^3)^ Deutsches Elektronen-Synchrotron DESY, Biomedical Research with X-rays (FS-BMX), Building 94, Room O4.003, Notkestraße 85, 22607 Hamburg, Germany

^4)^ Institute of Molecular Medicine, University of Lübeck, Ratzeburger Allee 160, Lübeck 23562, Germany

^5)^ Institute of Biology, Humboldt-Universität zu Berlin, Unter den Linden 6, Berlin 10099, Germany

**Table S1**: Crystallographic data for all fragment-bound structures after PanDDA quick refinement.

|  | MPro_F2XEntry_sg5_ B07 (7HUC) | MPro_F2XEntry_sg5_ B08 (7HUD) | MPro_F2XEntry_sg5_ D08 (7HUE) |
| --- | --- | --- | --- |
| Wavelength | 0.9184 | 0.9184 | 0.9184 |
| Resolution range | 21.84 - 1.52  (1.54 - 1.52) | 21.78 - 1.61  (1.63 - 1.61) | 20.64 - 1.7  (1.72 - 1.7) |
| Space group | C 1 2 1 | C 1 2 1 | C 1 2 1 |
| Unit cell | 113.4 53.09 44.41 90 102.58 90 | 112.55 52.81 44.69 90 102.65 90 | 112.35 53.06 44.81 90 102.21 90 |
| Total reflections | 224,178 (37,156) | 188,237 (30,990) | 159,285 (26,129) |
| Unique reflections | 39,298 (1,286) | 33,046 (1,045) | 28,400 (921) |
| Multiplicity | 5.7 (5.9) | 5.7 (5.9) | 5.6 (5.7) |
| Completeness (%) | 99.0 (97.7) | 99.4 (98.5) | 99.7 (98.9) |
| Mean I/sigma(I) | 12.9 (0.9) | 12.2 (0.9) | 13.6 (1.1) |
| Wilson B-factor | 24.5 | 28.1 | 30.1 |
| R-merge | 0.081 (1.773) | 0.087 (1.768) | 0.071 (1.53) |
| R-meas | 0.089 (1.944) | 0.096 (1.941) | 0.079 (1.684) |
| CC1/2 | 0.999 (0.486) | 0.999 (0.363) | 0.999 (0.481) |
| Reflections used in refinement | 39,298 (1,286) | 33,046 (1,045) | 28,400 (921) |
| Reflections used for R-free | 1,972 (142) | 1,658 (121) | 1,424 (104) |
| R-work | 0.203 | 0.199 | 0.198 |
| R-free | 0.235 | 0.229 | 0.233 |
| Number of non-hydrogen atoms | 2,932 | 2,500 | 2,515 |
| macromolecules | 2,784 | 2,399 | 2,417 |
| ligands | 38 | 22 | 26 |
| solvent | 110 | 79 | 72 |
| Protein residues | 306 | 306 | 306 |
| RMS(bonds) | 0.009 | 0.158 | 0.115 |
| RMS(angles) | 1.6 | 4.0 | 3.8 |
| Ramachandran favored (%) | 98.0 | 97.0 | 97.4 |
| Ramachandran allowed (%) | 1.6 | 2.6 | 2.3 |
| Ramachandran outliers (%) | 0.3 | 0.3 | 0.3 |
| Rotamer outliers (%) | 2.6 | 3.0 | 1.9 |
| Clashscore | 4.49 | 3.4 | 3.7 |
| Average B-factor | 27.1 | 30.3 | 33.7 |
| macromolecules | 27.1 | 30.2 | 33.6 |
| ligands | 26.3 | 38.1 | 39.5 |
| solvent | 28.3 | 31.0 | 35.1 |

|  | MPro_F2XEntry_ sg19_B05 (7I13) | MPro_F2XEntry_ sg19_B08 (7I14) | MPro_F2XEntry_ sg19_C02 (7I15) | MPro_F2XEntry_ sg19_C07 (7I16) |
| --- | --- | --- | --- | --- |
| Wavelength | 0.9184 | 0.9184 | 0.9184 | 0.9184 |
| Resolution range | 45.97 - 2.0 (2.05 - 2.0) | 46 - 1.95 (2 - 1.95) | 49.38 - 2.0  (2.05 - 2.0) | 45.74 - 2.13 (2.19 - 2.13) |
| Space group | P 21 21 21 | P 21 21 21 | P 21 21 21 | P 21 21 21 |
| Unit cell | 68.19 99.76 103.6 90 90 90 | 67.99 99.91 103.65 90 90 90 | 67.98 99.62 103.7 90 90 90 | 67.67 98.94 103.18 90 90 90 |
| Total reflections | 285,199 (47,097) | 297,835 (49,002) | 275,616 (46,960) | 222,723 (36,588) |
| Unique reflections | 48,132 (3,162) | 51,764 (3,422) | 48,179 (3,137) | 38,744 (2,730) |
| Multiplicity | 5.9 (6.1) | 5.8 (5.9) | 5.7 (6.1) | 5.8 (6.0) |
| Completeness (%) | 99.6 (99.8) | 99.8 (99.5) | 99.8 (99.7) | 98.4 (98.3) |
| Mean I/sigma(I) | 8.9 (1.3) | 6.8 (1.1) | 6.4 (1.1) | 6.8 (1.2) |
| Wilson B-factor | 30.2 | 31.0 | 28.7 | 34.4 |
| R-merge | 0.154 (1.33) | 0.143 (1.167) | 0.225 (1.444) | 0.201 (1.464) |
| R-meas | 0.168 (1.455) | 0.184 (1.525) | 0.247 (1.575) | 0.218 (1.583) |
| CC1/2 | 0.996 (0.496) | 0.994 (0.463) | 0.992 (0.408) | 0.992 (0.501) |
| Reflections used in refinement | 48,132 (3,162) | 51,764 (3,422) | 48,179 (3,137) | 38,744 (2,730) |
| Reflections used for R-free | 2,100 (138) | 2,100 (139) | 2,099 (136) | 1,938 (136) |
| R-work | 0.228 | 0.230 | 0.241 | 0.227 |
| R-free | 0.268 | 0.270 | 0.279 | 0.270 |
| Number of non-hydrogen atoms | 5,116 | 5,490 | 5,726 | 4,867 |
| macromolecules | 5,008 | 5,357 | 5,603 | 4,802 |
| ligands | 26 | 64 | 63 | 27 |
| solvent | 82 | 69 | 60 | 38 |
| Protein residues | 606 | 601 | 602 | 607 |
| RMS(bonds) | 0.014 | 0.014 | 0.014 | 0.013 |
| RMS(angles) | 1.7 | 1.7 | 1.7 | 1.7 |
| Ramachandran favored (%) | 95.9 | 96.8 | 96.7 | 95.5 |
| Ramachandran allowed (%) | 3.8 | 3.0 | 3.2 | 4.3 |
| Ramachandran outliers (%) | 0.3 | 0.2 | 0.2 | 0.2 |
| Rotamer outliers (%) | 2.3 | 1.8 | 4.3 | 2.2 |
| Clashscore | 2.6 | 5.9 | 4.4 | 4.6 |
| Average B-factor | 36.8 | 36.0 | 34.5 | 40.3 |
| macromolecules | 36.9 | 36.2 | 34.5 | 40.4 |
| ligands | 27.8 | 32.2 | 42.6 | 37.4 |
| solvent | 31.6 | 31.3 | 27.1 | 32.8 |

|  | MPro_F2XEntry_ sg19_C10 (7I17) | MPro_F2XEntry_ sg19_D04 (7I18) | MPro_F2XEntry_ sg19_D08 (7I19) | MPro_F2XEntry_ sg19_D11 (7I1A) |
| --- | --- | --- | --- | --- |
| Wavelength | 0.9184 | 0.9184 | 0.9184 | 0.9184 |
| Resolution range | 46 - 1.69  (1.73 - 1.69) | 45.89 - 2.0  (2.04 - 2.0) | 46.13 - 1.93  (1.97 - 1.93) | 49.2 - 1.73  (1.77 - 1.73) |
| Space group | P 21 21 21 | P 21 21 21 | P 21 21 21 | P 21 21 21 |
| Unit cell | 67.89 99.84 103.65 90 90 90 | 68.05 99.28 103.5 90 90 90 | 68.06 99.89 104.03 90 90 90 | 67.83 99.09 103.14 90 90 90 |
| Total reflections | 453,419 (75,927) | 277,043 (43,682) | 330,015 (51,853) | 430,503 (70,047) |
| Unique reflections | 78,658 (5,202) | 48,237 (3,121) | 53,905 (3,555) | 71,352 (4,667) |
| Multiplicity | 5.8 (6.0) | 5.7 (5.7) | 5.7 (5.6) | 6.0 (6.2) |
| Completeness (%) | 99.0 (99.3) | 99.9 (99.9) | 99.7 (99.8) | 97.5 (97.4) |
| Mean I/sigma(I) | 8.5 (0.9) | 9.0 (1.1) | 8.9 (1.2) | 12.2 (1.2) |
| Wilson B-factor | 24.8 | 31.4 | 29.9 | 24.0 |
| R-merge | 0.125 (1.647) | 0.154 (1.48) | 0.143 (1.167) | 0.104 (1.521) |
| R-meas | 0.137 (1.805) | 0.170 (1.631) | 0.158 (1.283) | 0.114 (1.66) |
| CC1/2 | 0.997 (0.333) | 0.997 (0.456) | 0.997 (0.495) | 0.999 (0.447) |
| Reflections used in refinement | 78,658 (5,202) | 48,237 (3,121) | 53,905 (3,555) | 71,352 (4,667) |
| Reflections used for R-free | 2,099 (139) | 2,099 (136) | 2,093 (138) | 2,101 (138) |
| R-work | 0.232 | 0.214 | 0.2307 | 0.212 |
| R-free | 0.253 | 0.254 | 0.2684 | 0.239 |
| Number of non-hydrogen atoms | 5,275 | 5,424 | 5,642 | 6,115 |
| macromolecules | 5,111 | 5,281 | 5,492 | 5,881 |
| ligands | 34 | 38 | 62 | 66 |
| solvent | 130 | 105 | 88 | 168 |
| Protein residues | 604 | 606 | 602 | 611 |
| RMS(bonds) | 0.015 | 0.014 | 0.013 | 0.014 |
| RMS(angles) | 1.8 | 1.7 | 1.6 | 1.6 |
| Ramachandran favored (%) | 97.2 | 97.0 | 97.7 | 97.7 |
| Ramachandran allowed (%) | 2.5 | 2.7 | 2.3 | 2.3 |
| Ramachandran outliers (%) | 0.3 | 0.3 | 0 | 0 |
| Rotamer outliers (%) | 3.0 | 2.5 | 3.4 | 2.7 |
| Clashscore | 4.2 | 3.6 | 3.7 | 5.4 |
| Average B-factor | 31.6 | 38.1 | 36.7 | 29.5 |
| macromolecules | 31.7 | 38.2 | 36.8 | 29.6 |
| ligands | 36.6 | 37.1 | 33.6 | 28.0 |
| solvent | 29.6 | 34.0 | 30.9 | 29.6 |

|  | MPro_F2XEntry_ sg19_E11 (7I1C) | MPro_F2XEntry_ sg19_F04 (7I1D) | MPro_F2XEntry_ sg19_G03 (7I1E) | MPro_F2XEntry_ sg19_G04 (7I1F) |
| --- | --- | --- | --- | --- |
| Wavelength | 0.9184 | 0.9184 | 0.9184 | 0.9184 |
| Resolution range | 40.19 - 1.95  (2 - 1.95) | 46.06 - 2.0  (2.05 - 2.0) | 45.98 - 1.88  (1.92 - 1.88) | 45.78 - 1.77  (1.81 - 1.77) |
| Space group | P 21 21 21 | P 21 21 21 | P 21 21 21 | P 21 21 21 |
| Unit cell | 68.11 99.56 103.94 90 90 90 | 67.96 99.47 103.93 90 90 90 | 67.92 99.59 103.67 90 90 90 | 67.69 99.33 103.18 90 90 90 |
| Total reflections | 307,300 (49,769) | 286,518 (47,203) | 333,013 (51,567) | 399,897 (64,014) |
| Unique reflections | 51,838 (3,403) | 48,092 (3,137) | 57,495 (3,746) | 68,396 (4,489) |
| Multiplicity | 5.9 (6.0) | 6.0 (6.2) | 5.8 (5.6) | 5.9 (5.9) |
| Completeness (%) | 99.6 (99.2) | 99.5 (99.9) | 99.3 (98.7) | 99.7 (99.6) |
| Mean I/sigma(I) | 10.1 (1.2) | 9.3 (1.1) | 8.6 (1.1) | 10.5 (1.0) |
| Wilson B-factor | 31.6 | 30.2 | 28.4 | 26.1 |
| R-merge | 0.126 (1.364) | 0.168 (1.61) | 0.142 (1.33) | 0.118 (1.668) |
| R-meas | 0.138 (1.492) | 0.184 (1.759) | 0.157 (1.466) | 0.130 (1.831) |
| CC1/2 | 0.997 (0.463) | 0.996 (0.380) | 0.997 (0.462) | 0.998 (0.420) |
| Reflections used in refinement | 51,838 (3,403) | 48,092 (3,137) | 57,495 (3,746) | 68,396 (4,489) |
| Reflections used for R-free | 2,100 (138) | 2,100 (137) | 2,099 (137) | 2,100 (137) |
| R-work | 0.225 | 0.227 | 0.219 | 0.220 |
| R-free | 0.251 | 0.268 | 0.248 | 0.259 |
| Number of non-hydrogen atoms | 5,080 | 5,962 | 5,030 | 5,432 |
| macromolecules | 4,966 | 5,857 | 4,903 | 5,249 |
| ligands | 28 | 38 | 27 | 38 |
| solvent | 86 | 67 | 100 | 145 |
| Protein residues | 602 | 601 | 605 | 602 |
| RMS(bonds) | 0.014 | 0.014 | 0.014 | 0.01 |
| RMS(angles) | 1.7 | 1.7 | 1.7 | 1.6 |
| Ramachandran favored (%) | 97.8 | 96.1 | 96.5 | 96.8 |
| Ramachandran allowed (%) | 2.2 | 3.4 | 3.3 | 3.0 |
| Ramachandran outliers (%) | 0 | 0.5 | 0.2 | 0.2 |
| Rotamer outliers (%) | 2.2 | 3.5 | 2.6 | 2.2 |
| Clashscore | 4.2 | 8.8 | 2.7 | 3.9 |
| Average B-factor | 38.1 | 36.2 | 35.9 | 33.5 |
| macromolecules | 38.2 | 36.2 | 36 | 33.5 |
| ligands | 29.5 | 49.9 | 35.9 | 44.2 |
| solvent | 32.5 | 28.2 | 30.5 | 31.2 |

|  | MPro_F2XEntry_ sg19_G09 (7I1G) | MPro_F2XEntry_ sg19_G10 (7I1H) | MPro_F2XEntry_ sg19_H03 (7I1I) | MPro_F2XEntry_ sg19_H11 (7I1J) |
| --- | --- | --- | --- | --- |
| Wavelength | 0.9184 | 0.9184 | 0.9184 | 0.9184 |
| Resolution range | 49.28 - 1.83  (1.87 - 1.83) | 46.68 - 1.98  (2.03 - 1.98) | 45.51 - 1.88  (1.92 - 1.88) | 49.47 - 1.85  (1.9 - 1.85) |
| Space group | P 21 21 21 | P 21 21 21 | P 21 21 21 | P 21 21 21 |
| Unit cell | 67.75 98.55 102.84 90 90 90 | 68.81 100.7 105.35 90 90 90 | 67.83 98.84 102.55 90 90 90 | 68.28 98.8 104.47 90 90 90 |
| Total reflections | 359,421 (55,530) | 293,691 (45,143) | 303,524 (40,576) | 352,058 (53,678) |
| Unique reflections | 61,199 (4,045) | 51,335 (3,378) | 55,928 (3,606) | 59,793 (3,949) |
| Multiplicity | 5.9 (5.7) | 5.7 (5.5) | 5.4 (4.7) | 5.9 (5.7) |
| Completeness (%) | 99.6 (100.0) | 99.8 (99.6) | 98.2 (96.7) | 98.8 (98.6) |
| Mean I/sigma(I) | 10.5 (1.2) | 8.7 (1.2) | 9.8 (1.0) | 7.8 (1.0) |
| Wilson B-factor | 28.3 | 30.9 | 30.5 | 27.8 |
| R-merge | 0.111 (1.426) | 0.143 (1.316) | 0.116 (1.448) | 0.151 (1.599) |
| R-meas | 0.122 (1.570) | 0.157 (1.454) | 0.128 (1.626) | 0.165 (1.760) |
| CC1/2 | 0.998 (0.460) | 0.996 (0.468) | 0.998 (0.389) | 0.996 (0.400) |
| Reflections used in refinement | 61,199 (4,045) | 51,335 (3,378) | 55,928 (3,606) | 59,793 (3,949) |
| Reflections used for R-free | 2,099 (138) | 2,099 (138) | 2,099 (135) | 2,101 (139) |
| R-work | 0.209 | 0.224 | 0.219 | 0.210 |
| R-free | 0.235 | 0.258 | 0.255 | 0.248 |
| Number of non-hydrogen atoms | 5,366 | 4,877 | 4,873 | 4,983 |
| macromolecules | 5,197 | 4,757 | 4,768 | 4,807 |
| ligands | 51 | 36 | 39 | 39 |
| solvent | 118 | 84 | 66 | 137 |
| Protein residues | 605 | 602 | 606 | 611 |
| RMS(bonds) | 0.014 | 0.01 | 0.014 | 0.009 |
| RMS(angles) | 1.7 | 1.7 | 1.7 | 1.5 |
| Ramachandran favored (%) | 97.3 | 97.3 | 97.5 | 97.9 |
| Ramachandran allowed (%) | 2.5 | 2.5 | 2.5 | 2.1 |
| Ramachandran outliers (%) | 0.2 | 0.2 | 0 | 0 |
| Rotamer outliers (%) | 2.4 | 1.3 | 1.7 | 1.1 |
| Clashscore | 4.5 | 1.6 | 3.0 | 3.0 |
| Average B-factor | 34.5 | 36.6 | 39.2 | 34.6 |
| macromolecules | 34.5 | 36.6 | 39.2 | 34.6 |
| ligands | 33.4 | 42.2 | 44.0 | 42.1 |
| solvent | 34.3 | 31.5 | 33.5 | 32.9 |

**Table S2**: Comparison between orthorhombic MPro crystals between Noske *et al.* (2021) and this study.

|  | Noske *et al.* | This study orthorhombic |
| --- | --- | --- |
| **Protein construct** | Immature MPro (50 amino acids per row)  **GAM**SGFRKMAFPSGKVEGCMVQVTCGTTTLNGLWLDDVVYCPRHVICTSE  DMLNPNYEDLLIRKSNHNFLVQAGNVQLRVIGHSMQNCVLKLKVDTANPK  TPKYKFVRIQPGQTFSVLACYNGSPSGVYQCAMRPNFTIKGSFLNGSCGS  VGFNIDYDCVSFCYMHHMELPTGVHAGTDLEGNFYGPFVDRQTAQAAGTD  TTITVNVLAWLYAAVINGDRWFLNRFTTTLNDFNLVAMKYNYEPLTQDHV  DILGPLSAQTGIAVLDMCASLKELLQNGMNGRTILGSALLEDEFTPFDVV  RQCSGVTFQ | Mature MPro (50 amino acids per row)  ---SGFRKMAFPSGKVEGCMVQVTCGTTTLNGLWLDDVVYCPRHVICTSE  DMLNPNYEDLLIRKSNHNFLVQAGNVQLRVIGHSMQNCVLKLKVDTANPK  TPKYKFVRIQPGQTFSVLACYNGSPSGVYQCAMRPNFTIKGSFLNGSCGS  VGFNIDYDCVSFCYMHHMELPTGVHAGTDLEGNFYGPFVDRQTAQAAGTD  TTITVNVLAWLYAAVINGDRWFLNRFTTTLNDFNLVAMKYNYEPLTQDHV  DILGPLSAQTGIAVLDMCASLKELLQNGMNGRTILGSALLEDEFTPFDVV  RQCSGVTFQ |
| **Crystallization condition** | 8% (w/v) PEG 4,000  0.1 M MES pH 6.7  5% (v/v) DMSO | 23.5% (w/v) PEG 1,500,  0.2 M MIB pH 7.7  5% (v/v) DMSO  1 mM TCEP  0.025 mM EDTA pH 7.0 |
| **Soaking condition** |  |  |
| Chemical composition | 8% (w/v) PEG 4,000  0.1 M MES pH 6.7  5% (v/v) DMSO  30% (w/v) PEG 400 | 23.5% (w/v) PEG 1,500  0.2 M MIB pH 7.7  5% (v/v) DMSO  1 mM DTT  0.025 mM EDTA pH 7.0 |
| Maximum fragment concentration | 40 mM | 100 mM |
| Soaking time | 4h | overnight |
| **Resolution** | 2.2 – 2.8 Å | Average of 1.94 Å |
